# Sex Differences in VEGF and PDGF Ligand and Receptor Protein Expression during Adipose Tissue Expansion and Regression

**DOI:** 10.1101/2025.02.14.638177

**Authors:** Yingye Fang, Silvia Gonzales-Nieves, Vincenza Cifarelli, P.I. Imoukhuede

## Abstract

Inadequate angiogenesis in obesogenic adipose tissue (AT) has been implicated in disrupted adipogenesis and metabolic disorders. Yet, key cellular and molecular regulators of AT angiogenesis remain largely unidentified. This study sought to identify the dysregulated elements within the Vascular Endothelial Growth Factor (VEGF) and Platelet-Derived Growth Factor (PDGF) systems during obesity progression. We employ a mouse model, comprising both male and female mice, to investigate the changes in the VEGF/PDGF concentration and their receptor distribution in AT during short- and long-term weight gain and weight loss. Our results reveal pronounced sex-specific differences in obesity progression, with male and female mice exhibiting distinct angiogenic ligand and receptor profiles under identical dietary interventions. This data also lays the groundwork for developing computational models of VEGF/PDGF signaling networks in AT, allowing for the simulation of complex biological interactions and the prediction of therapeutic outcomes.

## INTRODUCTION

Obesity affects over 650 million adults worldwide and remains a major global health concern, contributing to cardiometabolic disorders, chronic diseases, and certain cancers^1^. Studies in rodent models suggest that differences in adipose tissue (AT) angiogenesis during AT expansion may influence systemic metabolic outcomes ^2,3^. Notably, female mice on high-fat diet exhibit greater vascularity in gonadal white adipose tissue (gWAT) than males, which may contribute to healthier metabolic function in females^4^. However, the detailed regulation of angiogenic signaling proteins in AT during obesity progression, particularly with respect to sex-specific responses, is unclear.

The vascular endothelial growth factor (VEGF) family, comprising VEGF-A, VEGF-B, VEGF-C, VEGF-D, and placental growth factor (PlGF), are the primary regulators of angiogenic activity. These VEGFs share a highly conserved cystine-knot structure essential for receptor binding and activate receptor tyrosine kinases to mediate angiogenic signaling. The most well-characterized member of the VEGF family is VEGF-A, which acts through a family of cognate receptor kinases in endothelial cells to stimulate blood-vessel formation^5,6^. VEGF-A promotes the formation of capillaries within AT, ensuring adequate oxygen and nutrient supply, which is essential for AT expansion and metabolic activity^7^. Proper vascularization helps prevent AT low oxygen content, or hypoxia, and associated inflammation, both of which are linked to insulin resistance and metabolic dysfunction in human obesity^3^. Beyond its canonical roles in angiogenesis, VEGF-A is also as a critical regulator of AT energy metabolism by enhancing the delivery of glucose and lipids to adipocytes via improved blood flow and nutrient supply^8^. It also indirectly affects metabolic pathways by modulating AT inflammation and maintaining an anti-inflammatory microenvironment^9^. Overexpression of VEGF-A in AT is associated with increased vascularization but may also promote leaky vasculature, exacerbating inflammation and contributing to obesity-related metabolic disorders^10^. Conversely, insufficient VEGF-A signaling can result in poor vascularization, hypoxia, and chronic inflammation, further impairing AT function. Given its central role in regulating the interplay between vascular and metabolic systems, VEGF-A has been proposed as a promising therapeutic target for addressing metabolic diseases, including obesity, insulin resistance, and type 2 diabetes.

VEGF receptors, e.g., VEGFR1 and VEGFR2, are critical components of angiogenic signaling pathways and are broadly expressed in the adipose tissue^2^. VEGFR3 can regulate angiogenesis during early embryogenesis, but its primary function is as a lymphangiogenesis regulator^11^. VEGFRs share molecular structural features with Platelet-Derived Growth Factor Receptors (PDGFRs). VEGFRs and PDGFRs play complementary roles in vascular homeostasis. VEGFRs drive endothelial cell proliferation, migration, and survival, while PDGFRs, particularly PDGFRβ, are essential for the recruitment and stabilization of pericytes and smooth muscle cells, ensuring vascular integrity and preventing aberrant vessel formation^12^. We have previously identified binding across the VEGF and PDGF families using surface plasma resonance and predicted the significance of these cross-family binding interactions via computational modeling^13^. The cross-family PDGF binding was predicted to contribute up to 96% of VEGFR2 ligation in healthy conditions and cancers, introducing new mechanisms for VEGFR signaling. Thus, cross-family interaction between VEGF and PDGF signaling might regulate angiogenesis and vascular stability providing a novel layer of complexity in the angiogenic, vascular, and metabolic cues that regulate AT homeostasis.

Our goal was to investigate the dynamic expression patterns of VEGFRs, PDGFRs, and their ligands in the AT during diet-induced weight gain and regression weight loss in male and female mice. In pre-clinical models, female mice have an increased capacity for adipocyte enlargement in response to high-fat diet feeding, reduced AT inflammation, and lower fat deposition in the liver, which is associated with better metabolic health compared to male mice. ^14–16^ However, the VEGFRs and PDGFRs signaling profile remains unclear because, to date, preclinical studies investigating the VEGFRs and PDGFs signature in diet-induced obesity have been conducted mostly in male rodents. Results from our study reveal a distinct sex-specific signature for these angiogenic signaling proteins across key cell populations—endothelial cells, macrophages, and stromal cells—during AT remodeling in conditions of weight gain and weight loss. The data generated provides a foundation for building computational models of VEGF/PDGF signaling networks in AT, enabling the simulation of complex biological interactions and prediction of therapeutic outcomes.

## RESULTS

### Weight Changes of Male and Female Mice during HFD Feeding and HFD-to-LFD Dieting

To induce obesity, male and female C57BL/6J mice were challenged with high-fat diet (HFD) feeding for 12 weeks (short-term) or 24 weeks (long-term). Control groups were maintained on LFD for 12 or 24 weeks (**Figure 1A**). Gonadal white adipose tissue (gWAT) was collected from mice at Week 12 and Week 24 to assess gWAT mass, VEGF and PDGF ligand concentrations in gWAT, and receptor distribution on the surface of endothelial cells (ECs), macrophages, and stromal cells of gWAT (**Figure 1B**). Under HFD, the average body weight (BW) in male mice was 23.48 ± 0.28 g at week 0, increased to 47.33 ± 0.46 g at week 12, and further increased to 54.56 ± 0.86 g at week 24 (**Figure 1C**). The LFD control male BW was 22.19 ± 0.39 g at week 0, increased to 33.22 ± 0.57 g at week 12, and plateaued to 37.04 ± 1.2 g at week 24. A subset of HFD-fed male mice (BW: 46.5 ± 0.54 g) were switched to LFD at week 12; BW decreased to 35.4 ± 1.17 g at week 16, comparable to the control BW (35 ± 1.18 g). The average BW of the female group was 18.79 ± 0.22 g at week 0, increased to 30.08 ± 0.86 g on week 12, and reached 42.26 ± 3.01 g on week 24 (**Figure 1C**). The LFD control females started at 18.22 ± 0.34 g on week 0, increased BW to 23.75 ± 0.44 g on week 12, and plateaued at 24.2 ± 0.74 g on week 24. At week 12, a subset of HFD-fed female mice (BW: 31.58 ± 1.6 g) were switched to LFD; BW decreased to 23.56 ± 0.9 g on Week 14 and maintained similar BW until week 24 (24.78 ± 1.17 g) (**Figure 1C**). Overall, HFD and HFD-to-LFD dietary interventions effectively induced the progression and regression of obesity, respectively, in both sexes.

**Figure 1.**
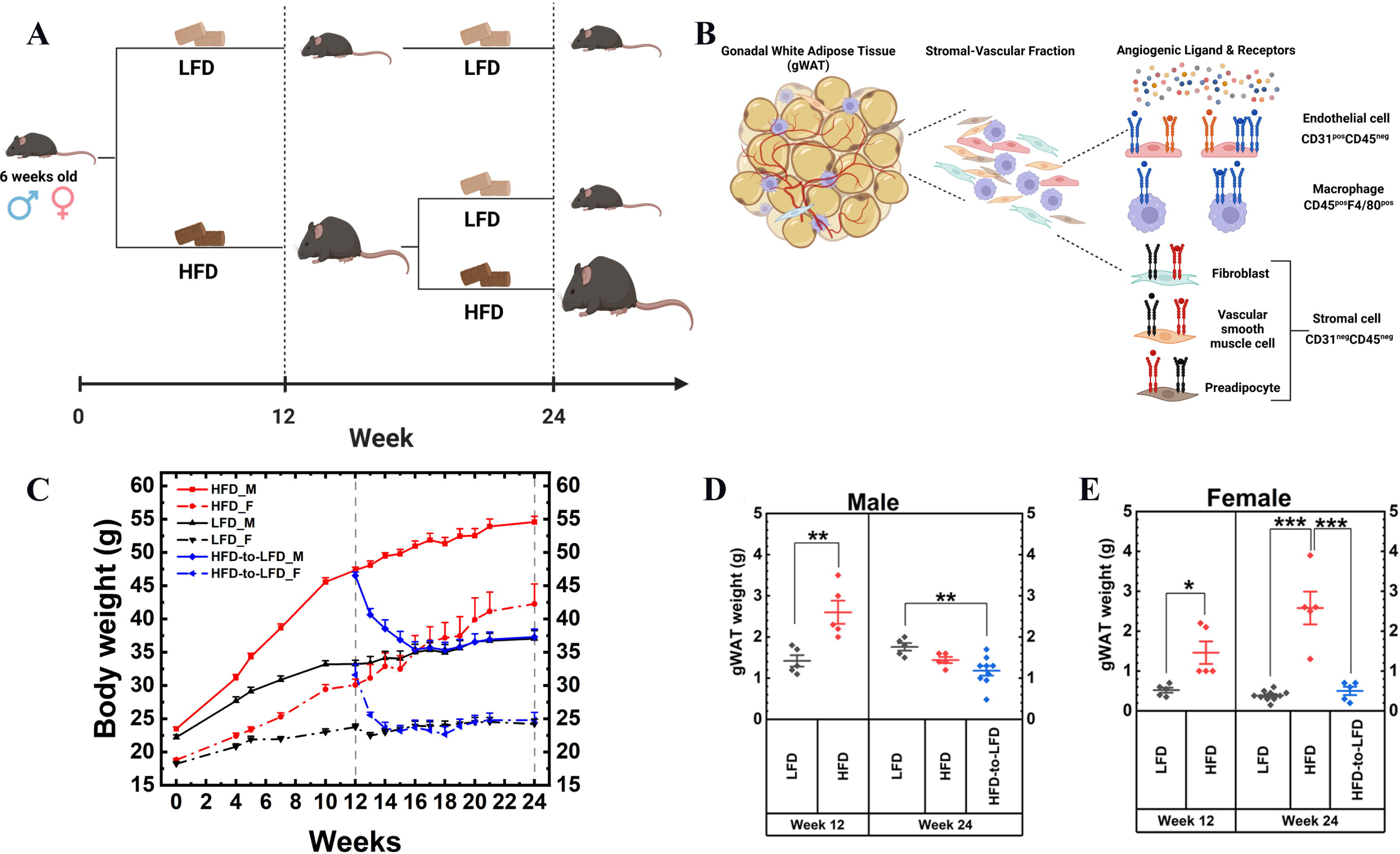
Experimental design overview. **(A)** Six-week-old male and female C57BL/6J mice underwent either 12-week (short-term) or 24-week (long-term) high-fat diet (HFD) feeding. A different cohort of male and female 12-week HFD-fed mice were switched to a low-fat diet (HFD-to-LFD) intervention to induce weight loss. Control groups were maintained on a low-fat diet (LFD) for 12 or 24 weeks. **(B)** Schematic summarizes measurements of angiogenic ligands (VEGF-A, VEGF-B, PLGF, PDGF-AA, PDGF-AB, and PDGF-BB in gonadal white adipose tissue (gWAT). Surface expression of Vascular Endothelial Growth Factor Receptors (VEGFRs) and Platelet-Derived Growth Factor Receptors (PDGFRs) was assessed in endothelial cells (CD31^positive^CD45^negative^F4/80^negative^) macrophages (CD31^negative^CD45^positive^F4/80^positive^) and stromal cells (CD31^negative^CD45^negative^F4/80^negative^) in gWAT stromal-vascular fraction (SVF) in the different dietary regimens as described in (A). **(C)** Body weight (g) in male and female mice during different dietary regimens. **(D-E)** Weight (g) of gWAT in male and female mice at the indicated time points/diet. All data represent the mean ± SEM. One-way ANOVA and post hoc Tukey test, * p < 0.05, ** p < 0.01, *** p < 0.001.

### Sex-specific Adipose Tissue Expansion during Long-Term HFD Feeding

Short-term weight gain resulted in a significant increase in gWAT mass from 1.42 ± 0.14 g to 2.6 ± 0.28 g in males (p = 0.005; **Figure 1D**) and from 0.52 ± 0.064 g to 1.46 ± 0.28 g in females (p = 0.01; **Figure 1E**), which was reversed to 1.18 ± 0.12 g (males) and 0.5 ± 0.1 g (females) by weight loss following the HFD-to-LFD dietary switch (**Figure 1D and 1E**). Sex differences in weight gain became evident after long-term HFD regimen (24 weeks), as gWAT mass in HFD-fed male mice was similar to the weights of LFD-fed mice at Week 24 (1.44 ± 0.07 g vs. 1.76 ± 0.09 g, p = 0.2; **Figure 1D**), whereas HFD-fed female mice maintained higher gWAT mass, compared to LFD-fed control (2.58 ± 0.41g vs. 0.39 ± 0.04 g, p < 0.0001; **Figure 1E**), indicating that gWAT lost the ability to expand in male mice after long-term HFD feeding, while female mice retained this capacity.

### Long-Term HFD Reduced VEGF-A in Male Mice but not in gWAT Female mice

VEGF-A is the primary pro-angiogenic growth factor. We measure VEGF-A levels in gWAT during short- and long-term HFD feeding and following the HFD-to-LFD dietary switch. We found that VEGF-A levels in gWAT were unchanged after short-term HFD feeding in both sexes (2.31 ± 0.38 g in HFD males and 4.22 ± 0.39 g in HFD females), compared to their LFD controls (2.31 ± 0.34 g in LFD males and 4.35 ± 0.8 g in LFD females) (**Figure 2A**). After long-term HFD feeding, female mice maintained the same VEGF-A levels as LFD control (7.43 ± 0.94 g vs. 7.55 ± 1.25 g), while long-term HFD males significantly reduced VEGF-A concentrations compared to their LFD counterparts (3.78 ± 0.26 g vs. 7.45 ± 0.65 g, p = 0.001; **Figure 2A**). Therefore, VEGF-A remained at baseline level during short-term weight gain and weight loss, but it significantly decreased in males during long-term weight gain.

**Figure 2.**
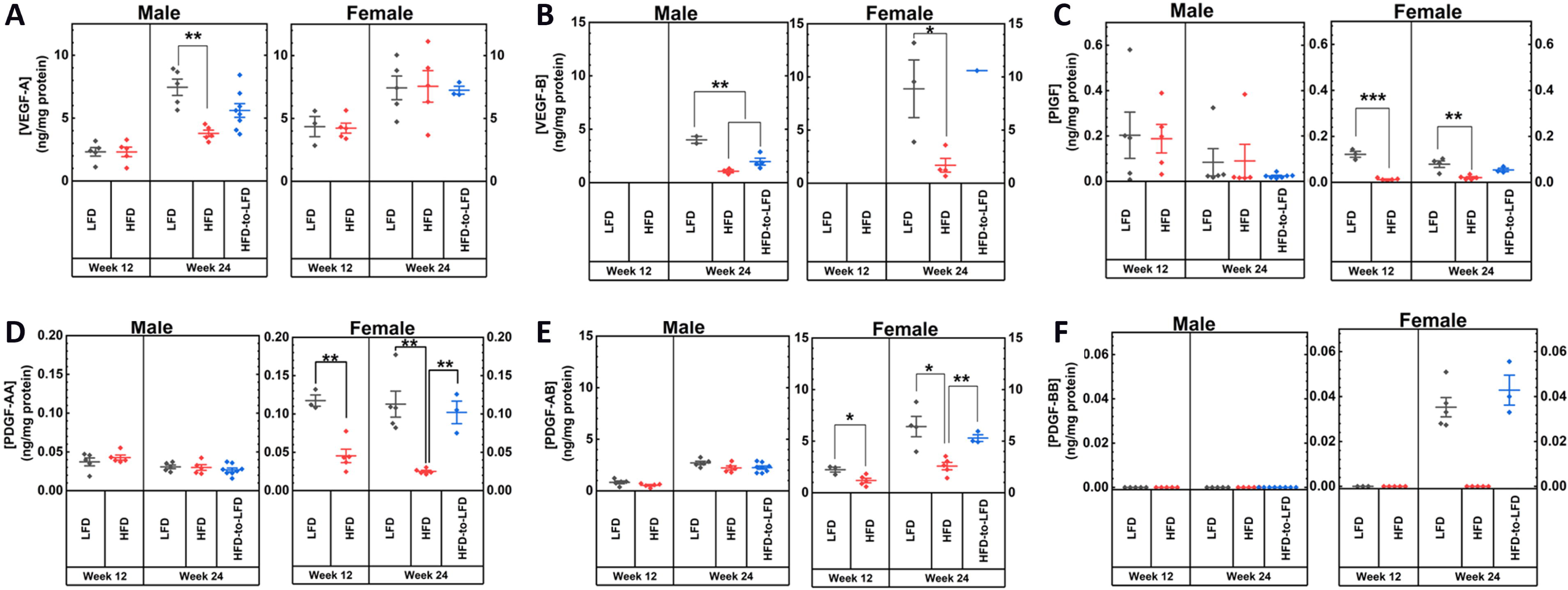
Measurements of angiogenic ligand concentrations in gWAT. ELISA was employed to measure the concentration of **(A)** VEGF-A, **(B)** VEGF-B, **(C)** PLGF, **(D)** PDGF-AA, **(E)** PDGF-AB, and **(F)** PDGF-BB. Tissue lysates (1 mg protein/ml solution) were used for the ELISA measurements. The ligand concentrations (ng ligands/ml solution) were normalized by the total protein in the solution (mg protein/ml solution) measured via the Bicinchoninic Acid (BCA) assay. All data represent the mean ± SEM. One-way ANOVA and post hoc Tukey test, * p < 0.05, ** p < 0.01, *** p < 0.001.

We next determined the cell source of VEGF-A by employing single-nucleus RNA sequencing (snRNA-seq). Cell types were stratified based on RNA marker expressions listed in **Suppl. Figure 1**. We compare *Vegfa* gene expressions across different gWAT populations namely adipocytes (*Adipoq*, *Pparg*, *Plin1*, *Lep*), endothelial cells (*Pecam1*, *Cdh5*, *Flt1*), and macrophages (*Adgre1*, *Lyz2*, *Ccl6*). Adipocytes were the main source of *Vegfa* (**Suppl. Figure 2**). We found that HFD feeding significantly decreased *Vegfa* RNA expression in adipocytes of male mice at Week 12 and 24, compared to LFD controls (**Suppl. Figure 3**). In contrast, HFD-fed female mice exhibited an increased *Vegfa* expression at Week 24 (**Suppl. Figure 3**). Overall, both protein and RNA measurements indicate that VEGF-A expression remained elevated in female mice but decreased in male mice following HFD feeding (**Figure 2A and Suppl. Figure 3**). VEGF-B, another member of the VEGF family that exclusively binds to VEGFR1^17^, was measured only in a subset of mice at Week 24 due to the limited tissue samples (**Figure 2B**). VEGF-B concentration was downregulated by HFD at Week 24 in both sexes (HFD male: 1.07 ± 0.15 g; HFD female: 1.69 ± 0.64 g), compared to their LFD controls (LFD male: 4.0 ± 0.32 g; LFD female: 8.87 ± 2.7g). The RNA expression of VEGF-B was also lower in HFD male and female mice, compared to their LFD controls, at Week 12 and Week 24 (**Suppl. Figure 3**).

### Female Mice Exhibited Higher Baseline Levels of PlGF, PDGF-AA, and PDGF-AB in gWAT

PlGF binds exclusively to VEGFR1 and plays a pro-angiogenic role in white AT expansion^18^. We found that short-term HFD did not change the PlGF concentration in male mice’s gWAT, compared to the LFD control (0.188 ± 0.06 vs. 0.20 ± 0.10 ng/mg protein), while it decreased female mice’s PlGF concentration (0.12 ± 0.012 vs. 0.011 ± 0.003 ng/mg protein, p = 0.00004) (**Figure 2C**). Similarly, long-term HFD did not change PlGF levels in males compared to LFD, but it increased PlGF levels in females (0.07 ± 0.03 vs. 0.02 ± 0.01ng/mg protein, p = 0.003) (**Figure 2C**). Thus, PlGF protein expression exhibited sex-specific patterns, where weight gain significantly decreased PlGF in female mice but did not affect male mice. Nevertheless, the RNA expression of PlGF by adipocytes was not altered by HFD feeding (**Suppl. Figure 3**).

Sex-specific expression patterns were also observed with PDGF-AA and PDGF-AB. Here, we show that, like PlGF, both PDGF-AA (**Figure 2D**) and PDGF-AB (**Figure 2E**) were at low levels in lean male mice (0.025-0.045 ng PDGF-AA/mg protein and 0.5-2.7 ng PDGF-AB/mg protein) but about threefold higher in lean females (0.11 ng PDGF-AA/mg protein and 5.3-6.4 ng PDGF-AB/mg protein). Notably, weight gain significantly reduced PDGF-AA and PDGF-AB levels in female mice, while male mice were unaffected. Furthermore, the HFD-to-LFD dietary switch reversed obesity-induced PDGF in female mice. The RNA expression of PDGF-A in adipocytes was unaltered by HFD feeding. PDGF-B was reduced by short-term HFD feeding in males; however, no difference was observed after long-term HFD feeding in either sex (**Suppl. Figure 3**).

PDGF-BB, another PDGF isoform, was detected at similar levels in lean female mice of both control and weight-loss conditions at Week 24 (∼ 0.04 ng/mg protein) (**Figure 2F**). However, PDGF-BB was not detected in sex- or diet-dependent conditions, suggesting that PDGF-BB concentrations may be below the test detection limit (< 0.03 ng/mg protein).

### Sex-Specific changes in SVF cell type composition During Weight Gain and Weight Loss

AT stromal-vascular fraction (SVF) consists of endothelial cells (ECs), macrophages, and a mixture of stromal cells, such as perivascular cells, fibroblasts, vascular smooth muscle cells, and preadipocytes^19^. Viable ECs were identified based on their Sytox Blue^neg^CD31^pos^CD45^neg^F4/80^neg^ phenotype; viable macrophages were Sytox Blue^neg^CD31^neg^CD45^pos^F4/80^pos^; and viable stromal cells were Sytox Blue^neg^CD31^neg^CD45^neg^F4/80^neg^ (**Figure 3A**). The surface expression levels of VEGFRs and PDGFRs were measured using PE-conjugated antibodies specific to each receptor on the respective cell types (e.g., PDGFRβ, **Figure 3B**). The percentages of ECs (**Figure 3C-D**) and macrophages (**Figure 3E-F**) in SVF are quantified in male and female mice via flow cytometry.

**Figure 3.**
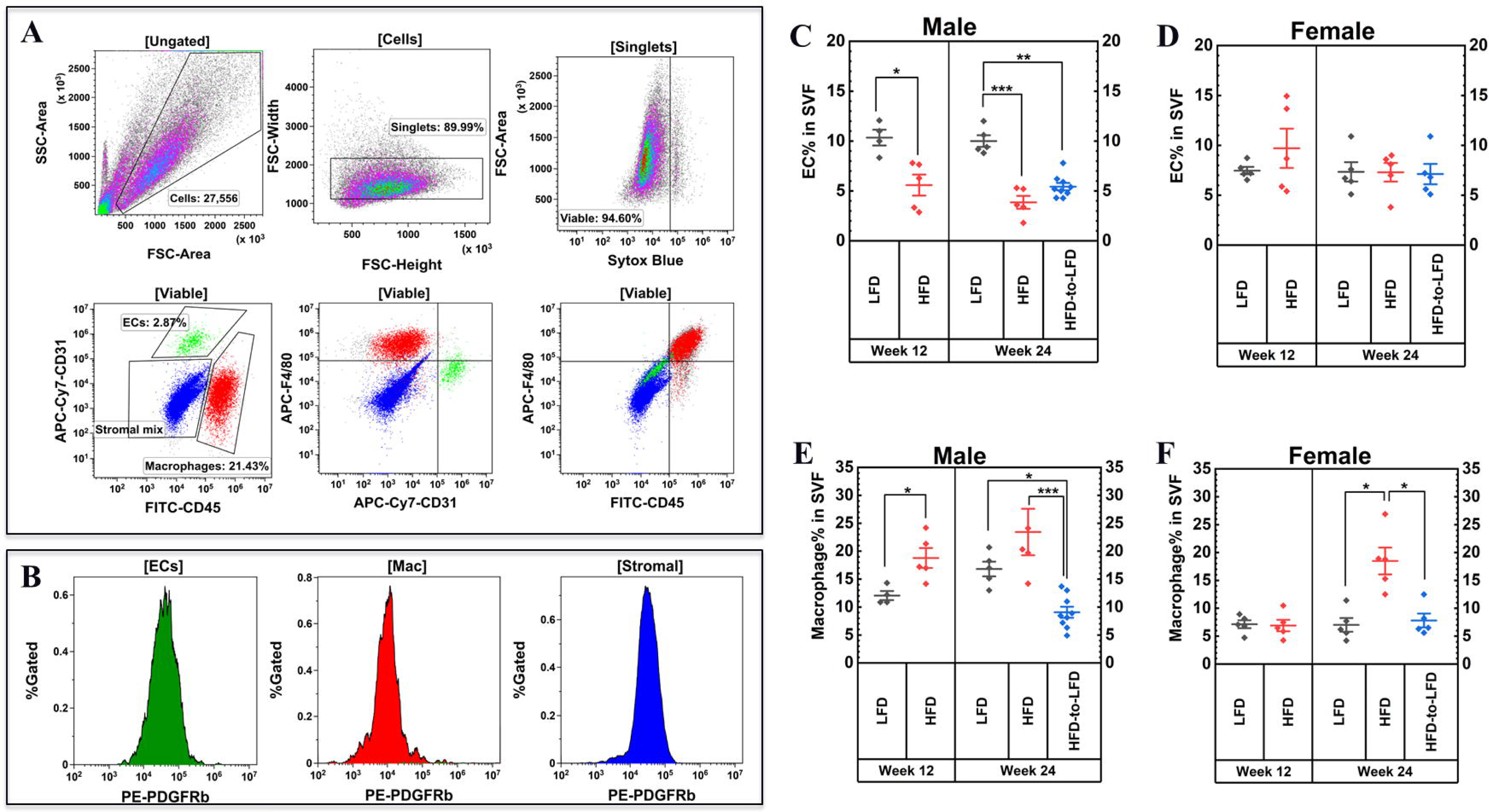
Flow cytometry analysis of the stromal vascular fraction (SVF) cells. **(A)** Flow cytometric cell strategy. Adipose SVF cells were dissociated into single cells, and stained with fluorophore-conjugated antibodies against CD31, CD45, F4/80, and one of the VEGFRs/PDGFRs. Debris, doublet cells, and dead cells were excluded as illustrated. The number and percentage of endothelial cells (CD31^positive^CD45^negative^F4/80^negative^), macrophages (CD31^negative^CD45^positive^F4/80^positive^), and stromal cells (CD31^negative^CD45^negative^F4/80^negative^) were assessed by flow cytometry. **(B)** PE signal distributions in ECs, macrophages, and stromal cells. **(C-D)** The percentage of endothelial cells (ECs) in the SVF. **(E-F)** The percentage of macrophages in the SVF. All data represent the mean ± SEM. One-way ANOVA and post hoc Tukey test, * p < 0.05, ** p < 0.01, *** p < 0.001.

Male mice showed decreased EC abundance after short- and long-term weight gain (expressed as EC% in SVF) (**Figure 3C**). The baseline EC% was 10% in LFD male control and 7.4% in female control. In males, the short-term weight gain decreased the EC% to 5.6% and the long-term weight loss further decreased it to 3.9%. The weight loss did not restore the male EC% to the control baseline (5.4% vs. 10%, p < 0.001). On the other hand, weight gain did not affect the EC% in female mice (**Figure 3D**).

The sex-specific differences in EC abundance observed in the flow cytometry data (SVF only) are consistent with the RNAseq analysis related to SVF and adipocytes. The RNAseq data revealed that the LFD male had three times more ECs than its short-term HFD counterpart (6.43% vs. 2.22%). Similarly, the long-term HFD-fed male exhibited a lower EC percentage (4.65%) compared to its LFD control (9.24%). In contrast, the difference between long-term LFD- and HFD-fed females was smaller (6.06% vs. 7.89%), with HFD-fed females showing a slightly higher percentage of ECs rather than a decrease (**Suppl. Figure 4**).

The percentage of macrophages (Sytox Blue^neg^CD31^neg^CD45^pos^F4/80^pos^ cells in SVF) was increased in male mice after both short- and long-term weight gain (**Figure 3E**). The macrophage% was 12-16% in the LFD male mice and 7-9% in the LFD female mice. The short-term weight gain increased the male macrophage% to 18.8% (p < 0.05) and the long-term weight gain further increased it to 23.4% (p < 0.05). Female mice did not show an increase in macrophage% after short-term weight gain but did exhibit a significant rise following long-term weight gain, from 7% to 18.5% (p < 0.01) (**Figure 3F**). Overall, weight gain led male mice to quickly decrease EC% and increase macrophage%, while female mice were more resistant to these adverse changes in vascular and immune cell composition.

### Endothelial VEGFR1 Concentrations Were Increased with Gonadal Adipose Tissue Size

We measured the abundance and surface distribution of VEGFR1, VEGFR2, VEGFR3, PDGFRα, and PDGFRβ in SVF cell populations (EC, macrophages, and stromal cells) using phycoerythrin (PE)-conjugated antibodies of the angiogenic receptors.

VEGFR1 was localized on the surface of ECs (**Figure 4A**) and macrophages (**Figure 5A**) but was absent on the stromal cells (**Figure 6A**) in the SVFs. On the ECs from the LFD and HFD-to-LFD (weight loss) groups, we quantified ∼2000 VEGFR1s/EC and considered the baseline VEGFR1 level of lean mice. The endothelial VEGFR1 concentrations significantly increased after short-term weight gain in both male (to 4706 VEGFR1s/EC, p = 0.033) and female mice (3372 VEGFR1s/EC, p = 0.017) (**Figure 4A, Week 12**). The long-term HFD maintained the elevated endothelial VEGFR1 concentration in female mice (4993 VEGFR1s/EC). On the contrary, long-term HFD male mice had lower endothelial VEGFR1 concentration when compared to that of short-term HFD male mice (2271 vs. 4706 VEGFR1/EC, p = 0.048, **Figure 4A, Week 24**). Notably, the upregulation and downregulation of endothelial VEGFR1 levels act in concert with gWAT expansion and shrinkage in HFD-fed male and female mice (**Figure 1D-E**), suggesting that endothelial VEGFR1 may play a regulatory role in gWAT expansion. However, VEGFR1 RNA (Flt-1) expression levels in ECs were unaltered by diets (**Suppl. Figure 5**).

**Figure 4.**
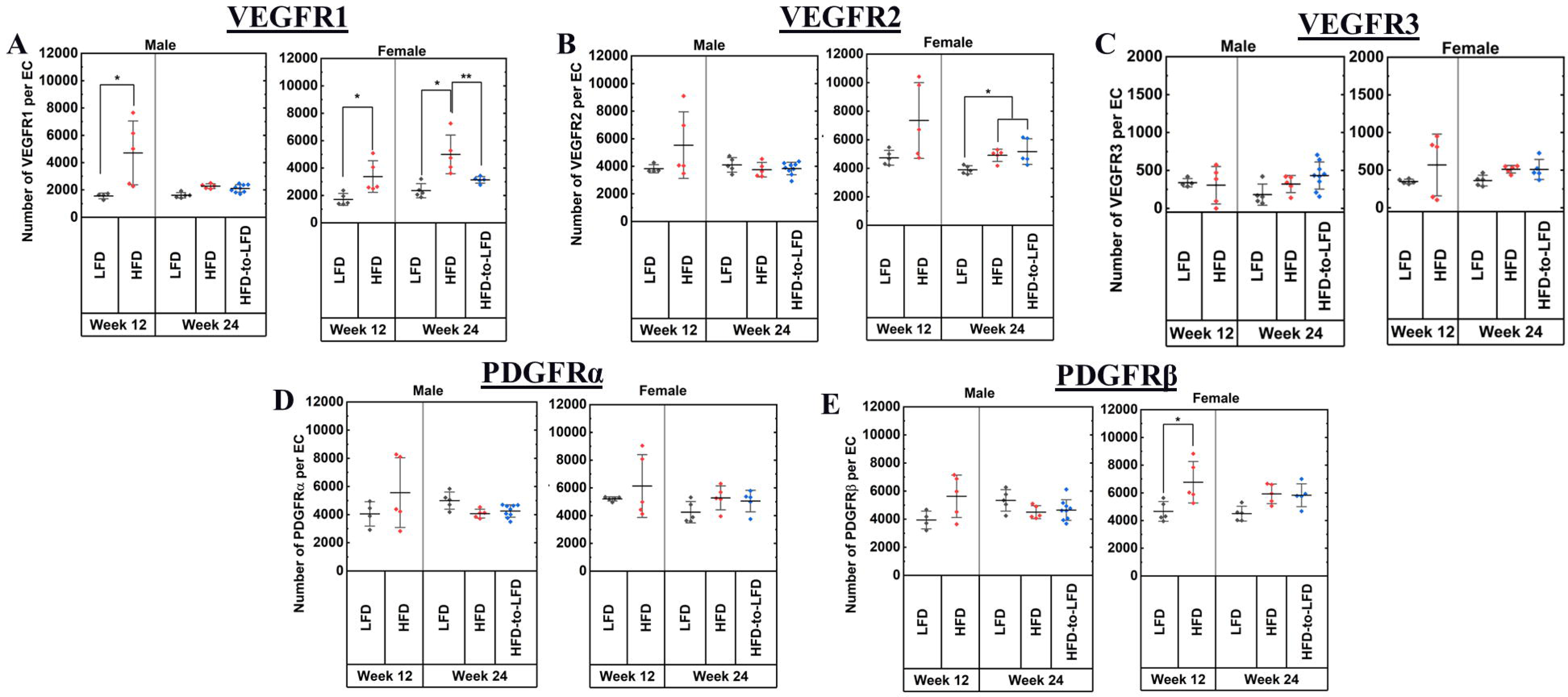
VEGFR1, VEGFR2, VEGFR3, PDGFRα, and PDGFRβ concentrations on the surface of endothelial cells. Quantitative flow cytometry was employed to measure endothelial cell surface (CD31^positive^CD45^negative^F4/80^negative^) surface expression of **(A)** VEGFR1 **(B)** VEGFR2, **(C)** VEGFR3, (**D)** PDGFRα and (**E**) PDGFRβ in male and female mice at the indicated time points/diet regimens. All data represent the mean ± SEM. One-way ANOVA and post hoc Tukey test, * p < 0.05, ** p < 0.01, *** p < 0.001.

**Figure 5.**
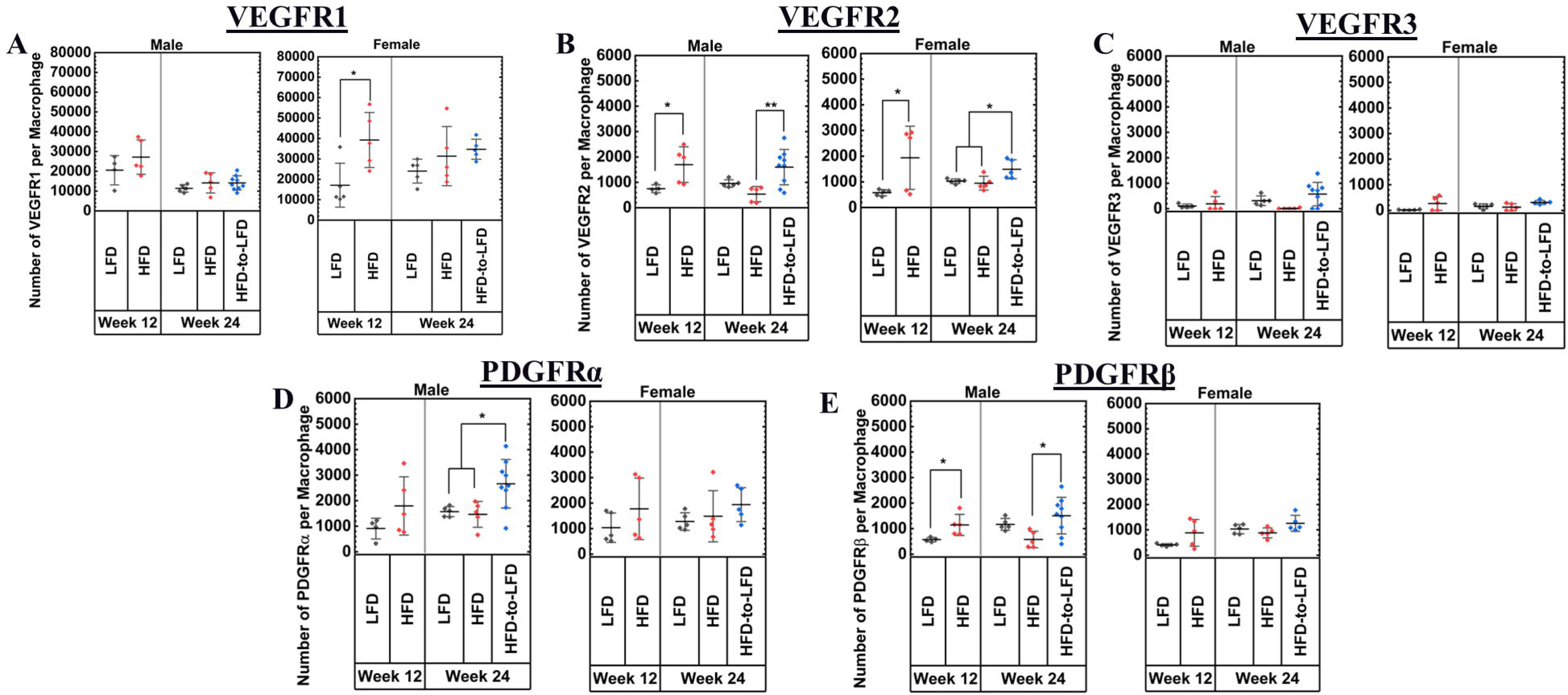
VEGFR1, VEGFR2, VEGFR3, PDGFRα, and PDGFRβ concentrations on the surface of macrophages. Quantitative flow cytometry was employed to determine macrophage (CD31^negative^CD45^positive^F4/80^positive^) surface expression of **(A)** VEGFR1 **(B)** VEGFR2, **(C)** VEGFR3, **(D)** PDGFRα and (**E**) PDGFRβ in male and female mice at the indicated time points/diet regimens. All data represent the mean ± SEM. One-way ANOVA and post hoc Tukey test, * p < 0.05, ** p < 0.01, *** p < 0.001.

**Figure 6.**
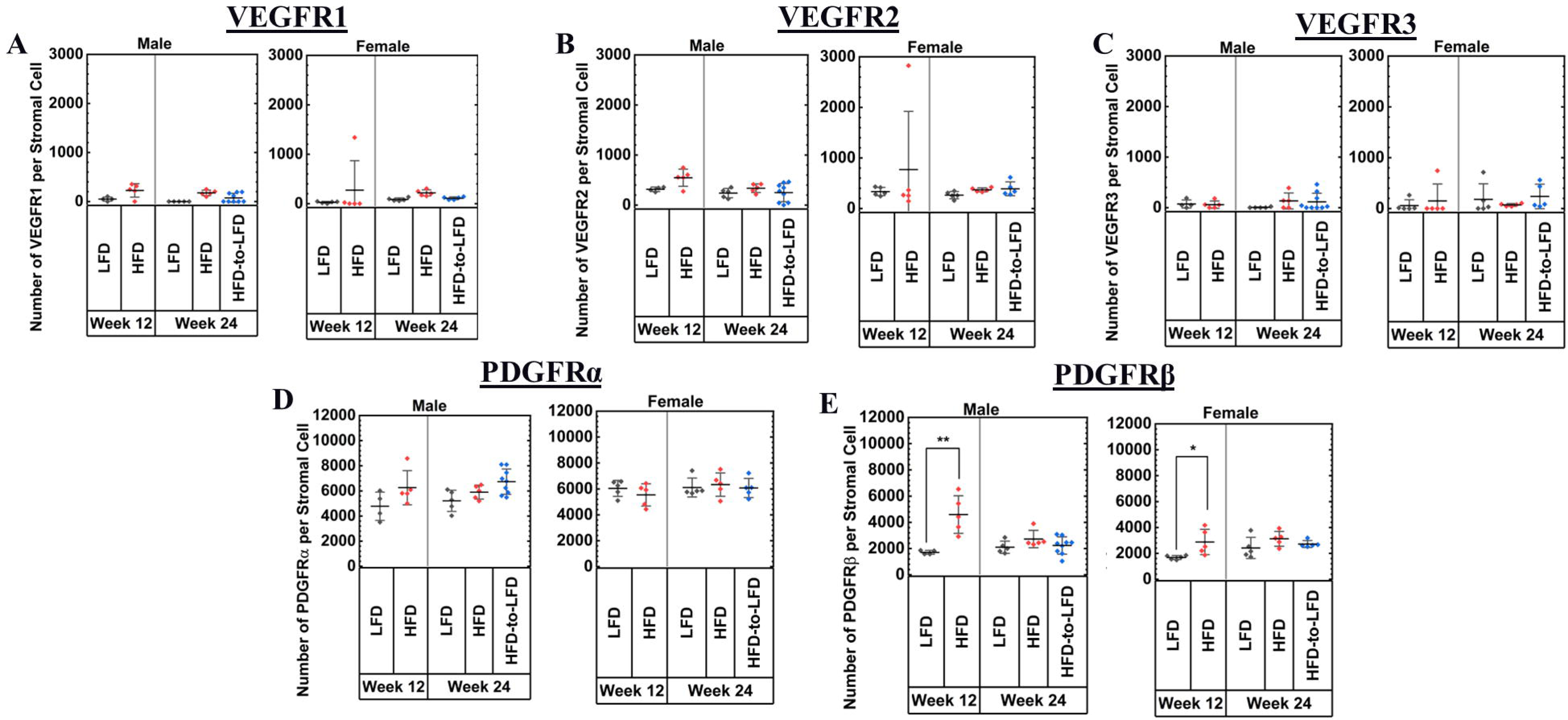
VEGFR1, VEGFR2, VEGFR3, PDGFRα, and PDGFRβ concentrations on the surface of stromal cells. Quantitative flow cytometry was employed to measure levels of **(A-C)** VEGFR1, VEGFR2, and VEGFR3, **(D)** PDGFRα and (**E**) PDGFRβ, in male and female mice at the indicated time points/diet regimens. All data represent the mean ± SEM. One-way ANOVA and post hoc Tukey test, * p < 0.05, ** p < 0.01, *** p < 0.001.

### Macrophage VEGFR1 Concentrations Were Transiently Elevated in Females After Short-Term Weight Gain

We found that gWAT macrophages have approximately ten times more VEGFR1 than gWAT ECs (**Figure 5A**). The short-term HFD significantly increased macrophage VEGFR1 in female mice, from 17,084 to 39200 VEGFR1s/cell (p = 0.02). On the other hand, the increase in VEGFR1 expression during short-term HFD male mice was not statistically significant, ranging from 20,540 to 27,190 VEGFR1s/cell (p = 0.26). We did not observe differences in VEGFR1 levels in long-term HFD. Thus, the sex difference in macrophage VEGFR1 was present at the early stage of weight gain but not after long-term weight gain.

### Macrophage VEGFR2 Concentrations Were Transiently Elevated After Short-Term Weight Gain

VEGFR2 is the most prominent pro-angiogenic receptor in the VEGFs/VEGFRs system that regulates the angiogenic hallmarks, including EC proliferation and migration^20^. We found VEGFR2 to be predominately expressed on the surface of ECs, ranging between 3817 and 7344 VEGFR2 per EC (**Figure 4B**), between 750 and 1930 VEGFR2s/macrophage (**Figure 5B**), and under the detection threshold of 400 receptors/cell for stromal cells among the different groups (**Figure 6B**).

Endothelial VEGFR2 concentrations did not significantly vary among the different diet groups in males (**Figure 4B**), while the long-term LFD females had lower endothelial VEGFR2 levels compared with their HFD and HFD-to-LFD counterparts (3885 vs. 4906 VEGFR2s/EC, p = 0.014; 3885 vs. 5168 VEGFR2s/EC, p = 0.048) (**Figure 4B**).

Elevated VEGFR2 protein expression has been observed in M2 macrophages, commonly associated with the pro-angiogenic phenotype, in contrast to the pro-inflammatory M1 phenotype.^21^ We show that AT macrophages exhibited < 1000 VEGFR2 per cell in lean mice (LFD). After short-term HFD, VEGFR2 levels doubled in male mice (p = 0.03) and tripled in female mice (p = 0.039) (**Figure 5B**). This difference was not observed at the RNA level, suggesting that the changes in protein levels may be attributed to post-translational regulation (**Suppl. Figure 5**). After long-term HFD, the differences between LFD and HFD diminished in both male and female mice, suggesting a shift in macrophage phenotype away from a pro-angiogenic M2 state with prolonged weight gain. VEGFR3 is expressed in endothelial cells where it acts as an important signaling mediator primarily for lymphangiogenesis^22^ and angiogenesis^23^. VEGFR3 was not detected on the surface of vascular ECs (**Figure 4C**), macrophages (**Figure 5C**), or stromal cells in AT (**Figure 6C**).

### PDGFRβ Concentrations Were Elevated in EC, Macrophage, and Stromal Cells in a Sex-specific Manner

PDGFRβ concentrations were significantly upregulated by HFD in female mice, whereas PDGFRα concentrations remained relatively stable across diet groups over time. Specifically, short-term HFD increased endothelial PDGFRβ concentration in female mice, from 4664 to 6770 PDGFRβs/EC (p = 0.031), but not in males (**Figure 4E**). In contrast, short-term HFD increased macrophage PDGFRβ concentrations in males from 570 to 1145 PDGFRβs/cell (p = 0.031), but not in females (p = 0.079) (**Figure 5E**). However, after long-term HFD, the differences between LFD and HFD in males were diminished. Nevertheless, the PDGFRβ RNA (Pdgfrb) expression by macrophages was significantly higher in long-term HFD males than its LFD control (**Suppl. Figure 6**).

Stromal PDGFRβ concentrations were increased after short-term HFD in both males (p = 0.0055) and females (p = 0.027), while this increase diminished after long-term weight gain in both sexes (**Figure 6E**). These data together highlight a sex-specific PDGFRβ upregulation in response to weight gain in the three main SVF cell types.

## DISCUSSION

Our study quantifies the expression of angiogenic proteins in gWAT during different dietary feeding regimens and highlights three key differences between male and female mice: (1) Short-term HFD feeding doubled the expression of endothelial VEGFR1 levels in both sexes; after long-term HFD feeding, the VEGFR1 level remained elevated in females but diminished in males. (2) Long-term HFD-fed males had lower VEGF-A in gWATs than lean males, whereas HFD-fed females and lean females had the same VEGF-A levels. Long-term HFD feeding reduced the VEGF-B levels in both sexes. (3) Levels of PDGF-AA and PDGF-BB in males were low and remained unaffected by HFD feeding. Compared to lean males, lean females had about three times higher baseline levels of PDGF-AA and -BB, which significantly decreased after short-and long-term HFD feeding. These findings identify the differentially regulated angiogenic signaling ligands and receptors that may contribute to previously reported higher vascularity^4^ and lower inflammation^24^ in female mice during HFD feeding. Aligned with previous studies, our obese female mice exhibited sustained AT expandability and higher vascular cell presence (i.e., EC%) in the AT, compared to their male counterparts after 24 weeks of HFD feeding.^14,24,25^ Our study aimed to quantitatively characterize angiogenic signaling proteins, as they may play a crucial role in regulating AT expansion, vascular cell abundance, and related metabolic functions in obese mice.

PlGF is a VEGF family member that exclusively binds to and signals through VEGFR1^18^. In non-obesogenic AT, PlGF has minimal influence, as neither its overexpression nor deletion has any discernible impact on the vascular or metabolic well-being of animals under LFD. ^26–28^ However, in obesogenic AT, PlGF^-/-^ male^26^ and PlGF^-/-^ female ^27^ mice displayed more severe metabolic disorders due to impaired vascularization, compared to their wildtype controls, suggesting the critical role of PlGF in AT vascular health in obesity. On the other hand, PlGF-overexpressing male mice also developed more severe metabolic disorders than wildtype due to elevated inflammation within obesogenic AT.^28^ These findings suggest that balanced PlGF expression is essential to prevent adverse effects on both vascularization and inflammation in AT during obesity.

Our data show that PlGF protein concentrations were reduced in HFD-fed female mice compared to their LFD controls, while PlGF levels in HFD-fed male mice remained unchanged. Notably, weight loss restored PlGF levels in female mice, suggesting that the weight gain-associated reduction in PlGF is reversible with dietary intervention. Our results may appear contradictory to a previous study that failed to detect PlGF protein in gWAT using ELISA ^18^. We believe that this discrepancy is due to test sensitivity (6 ng/mg protein). In our study, we used an ELISA assay with a higher sensitivity of 0.001 ng/mL (equivalent to 0.001 ng/mg protein when 1 mg of total protein is dissolved in 1 mL solution). With this method, we measured PlGF concentrations in gWAT ranging from 0 to 0.6 ng/mg protein.

Like PlGF, PDGF-AA and PDGF-AB were downregulated within obesogenic AT in a female-specific, reversible manner, suggesting a synergistic mechanism among these three ligands in regulating AT angiogenesis molecular signature. We also measured PDGF-BB, which induces the detachment of PDGFRβ^+^ perivascular cells from blood vessels, allowing vascular sprouting that increases adipose vascularity.^29^ However, we detected PDGF-BB only in the control and weight loss female mice at Week 24. To provide context to these results, PDGFs act in concert with proangiogenic factors, such as VEGF-A and PlGF, to induce the formation and stabilization of blood vessels.^30^ PDGF-AA, PDGF-BB, and PDGF-AB all bind to PDGFRα, while PDGF-BB and PDGF-AB additionally bind to PDGFRα. Moreover, PDGFs may activate VEGFR signaling because they demonstrate a stronger binding affinity for VEGFR2 compared to their binding to PDGFRs.^13^ The inability to detect PDGF-BB levels may be attributed to the possibility that the PDGF-BB level is less than 0.03 ng/mg protein, which falls below the detection limit of the ELISA kit. Overall, PlGF, PDGF-AA, and PDGF-AB were always downregulated in HFD female mice and remained unchanged in HFD male mice. This discovery points to the presence of a notable sex-specific PDGF signaling mechanism in adipose tissue, warranting further investigation.

VEGF-A concentrations in gWAT were not affected by short-term HFD feeding. Long-term HFD feeding (24 weeks) decreased VEGF-A concentration in males, while levels in females remained similar to their LFD control. These observations align with previous studies indicating that female mice have higher levels of pro-angiogenic factors in their adipose tissue compared to males^4^. Our snRNA-seq data suggested that the reduced VEGF-A protein concentrations in male mice are due to a decrease in its RNA expression by adipocytes because adipocytes are the main source of VEGF-A in adipose tissue^31^. Our observations aligned with a qPCR study, which found that female mice displayed greater levels of *Vegfa* mRNA after 16 weeks of HFD feeding, compared to their male counterparts.^4^ Another potential mechanism for the VEGF-A reduction in male mice could be adipocyte loss following long-term HFD feeding, which may exacerbate impaired angiogenesis in adipose tissue.^8,32^

VEGF-B contributes to adipose tissue angiogenesis. Robciuc et al.^33^ showed that VEGF-B binding to VEGFR1 promotes angiogenesis via the VEGF/VEGFR2 pathway. We show that VEGF-B is downregulated by HFD in both sexes at Week 24, likely due to a decrease in VEGF-B RNA (*vegfb*) expression by adipocytes, as suggested by our snRNA-seq data. Therefore, the impaired gWAT expansion observed in male mice at Week 24 may result from a combination of decreased VEGF-B, male-specific reduction of VEGFR1 on endothelial cells, and male-specific reduction of VEGF-A levels.

We further highlight our discovery of remarkable changes in VEGFR1 during obesity progression. The observed changes in VEGFR1 protein levels were likely driven by post-translational processes, such as the trafficking of VEGFR1 from intracellular compartments to the cell surface, because our snRNA-seq data suggests that HFD feeding does not alter VEGFR1 RNA (Flt-1) expression in gWAT endothelial cells. A crucial switch that controls angiogenesis is the balance between the VEGFR1 and VEGFR2 levels on the surface of endothelial cells. Typically, angiogenesis remains mostly inactive in adults, unless prompted by a physiological demand namely menstruation^34^, adipose tissue expansion^7^, and wound healing^35^, or triggered by a pathological stimulus (e.g., tumor growth^36^). Reflecting this inactivity, the endothelial cells within non-obesogenic AT, have ∼2000 VEGFR1 and ∼4000 VEGFR2 per cell. Similar VEGFR levels were observed in *in vitro* human endothelial cell lines when cultured at a low VEGF concentration (2 pM)^37^. When culturing those cell lines at a high VEGF concentration (1 nM), VEGFR1 levels can double or even triple^37^. VEGFR1 has also been shown to be increased on ECs from ischemic hindlimb when reperfusion rates to the hindlimb are greatest. This study shows that, in male mice, the endothelial VEGFR1 level doubled after short-term HFD as their AT expanded but reduced to the ‘inactive’ level after long-term HFD, coinciding with the loss of AT expandability in the MUO male mice.

The upregulation of VEGFR1, but not of VEGFR2 and VEGFR3 on the surface of endothelial cells during active adipose tissue expansion suggests that VEGFR1 might play a critical role in regulating this process. Traditionally, VEGFR1 has been considered a decoy receptor that sequesters VEGF-A, preventing its interaction with VEGFR2. However, VEGFR1 upregulation also occurs in pathological conditions, such as hypoxia and cancers^38,39^. It is possible that VEGFR1 upregulation may facilitate gWAT expansion, potentially by promoting endothelial cell proliferation and migration. Supporting this hypothesis, our data show that male mice exhibited reduced EC abundance in gWAT following HFD feeding, as evidenced by flow cytometry analysis of the SVF and snRNA-seq data of the gWAT, whereas females did not. Reduction in EC abundance in male mice was accompanied by reduced gWAT expandability, while females maintained gWAT expansion during long-term HFD feeding. Notably, during long-term HFD feeding, VEGFR1 protein expression on endothelial cells doubled in female gWAT compared to their LFD controls, whereas males showed no significant change relative to their LFD controls. These findings suggest that elevated VEGFR1 on ECs may influence endothelial function and tissue remodeling, contributing to the sex-specific differences in EC abundance and gWAT expansion observed during HFD exposure.

Regarding proliferation, high VEGFR1 is found on highly proliferative ‘stalk’ cells that stabilize angiogenic sprouts, while high VEGFR2 is found on ‘tip’ cells that form angiogenic sprouts.^40^ Under obesogenic conditions, our female mice exhibited persistently elevated VEGFR1 on ECs, but not in macrophages or stromal cells suggesting ECs in female mice are the proliferative ‘stalk’ type. On the other hand, elevated endothelial VEGFR1 in male mice was diminished after long-term weight gain when their AT failed to expand. Moreover, the abundance of ECs was preserved in HFD-fed female mice but significantly decreased in HFD-fed males, which further supports the higher proliferative potential of female adipose endothelial cells during HFD feeding. Our findings align with previous studies reporting increased blood vessel formation in female mice, driven by enhanced vascular proliferation processes.^24^

Regarding migration, VEGFR1 signaling promotes migration of tumor cells^41^, myeloid cells^42,43^, and macrophages^44^. VEGFR1 promotes RAW264.7 macrophage migration through PLCg and PI3K signaling pathways^41^. During long-term HFD feeding, pro-inflammatory macrophages are increasingly recruited to the visceral AT^45^, which leads to chronic inflammation and associated metabolic diseases. We show that macrophage VEGFR1 level was transiently elevated after short-term HFD in female mice, but not in male mice. However, the elevated macrophage VEGFR1 level did not increase macrophage recruitment in female mice. Conversely, male mice on short-term HFD did not show a significant rise in macrophage VEGFR1 but did exhibit increased macrophage presence in the gWAT. Long-term HFD diminished the elevated macrophage VEGFR1 in obese female mice, yet both sexes displayed significantly higher macrophage presence, which further contradicts the expectation that high VEGFR1 levels promote macrophage recruitment. The role of elevated macrophage VEGFR1 in gWAT remodeling in female mice requires further investigation because understanding this mechanism could provide important insights into the sex-specific regulation of macrophage dynamics and their role in obesity-related inflammation.

However, the migratory role of VEGFR1 signaling in human endothelial cells has not been established, potentially due to the low abundance of surface VEGFR1, ranging from a few hundred to 1800 VEGFR1 molecules per cell on the ‘inactive’ endothelial cells *in vitro* and *ex vivo* ^37,46,47^. Our study shows that lean mice had ∼2000 VEGFR1 per endothelial cell, reflecting ‘inactive’ angiogenesis. In contrast, obese mice with expanding adipose tissue exhibited doubled endothelial VEGFR1 levels, which may be critical for promoting cell migration to meet angiogenic needs during adipose tissue expansion. Thus, endothelial VEGFR1 and its signaling pathway in adipose tissue may be targeted to promote cell proliferation and migration, enhancing AT vascularization and expandability in obesogenic conditions.

VEGFR1 also exists in a soluble form, known as sVEGFR1 (or sFlt-1), which binds to VEGF-A, VEGF-B, and PlGF, thereby inhibiting their interactions with membrane-bound VEGFRs and modulating angiogenesis. Nevertheless, Tahergorabi and Khazaei reported that HFD did not significantly alter serum sVEGFR1 concentrations in male C57BL/6 mice^48^. Currently, there is limited research on how HFD affects sVEGFR1 expression in adipose tissue, indicating a need for further investigation in this area.

PDGFRα and PDGFRβ are predominantly expressed by stromal cells, such as fibroblasts, pericytes, and tissue progenitor cells^49^. High PDGFR expressions are also found in endothelial cells in certain tissue environments, such as bone^50^ and tumor^51^. This study demonstrates that PDGFRs are present on the surface of endothelial cells in gonadal adipose tissue, suggesting a direct role for PDGFR signaling in regulating adipose angiogenesis.

Stromal cells exhibit similar PDGFRα concentrations across all sex-diet conditions. Stromal cells, consisting of pericytes, preadipocytes, and fibroblasts, support angiogenesis likely through PDGFRs because they do not have VEGFRs on the cell surface. This observation aligns with the understanding that adult PDGFRα+ cells do not significantly contribute to adipogenesis, and deleting *Pdgfra* gene in adult adipose lineage did not affect AT homeostasis.^52^ On the other hand, increased *Pdgfrb* mRNA was previously reported in 12-week HFD male mice, and here we show increased PDGFRβ protein levels in both male and female mice after 12-week HFD. The increased *Pdgfrb* mobilized PDGFRβ^+^ pericytes to detach from the blood vessels in AT, enabling vascular sprouting in response to angiogenic demand.^29^ The same study demonstrated the significant role of stromal PDGFRβ in adipose angiogenesis with PDGFRβ-KO HFD mice, which had a normal number of pericyte-covered vessels but reduced AT vascularization, compared to the wild-type HFD mice. ^29^ Notably, we show that the increase in PDGFRβ protein expression was diminished by long-term HFD. Therefore, the transient upregulation of stromal PDGFRβ may be critical for increasing vascularity during early AT expansion, suggesting that the prime window for stromal PDGFRβ to promote angiogenesis is during the early stages of obesity development.

Similar patterns of PDGFR expression were observed in adipose ECs. Surface PDGFRs are typically undetectable in quiescent endothelial cells, while ∼3000 PDGFRβs/EC were detected during vascular tubule formation in the co-culture of endothelial cells and fibroblasts.^53^ PDGFRs are also highly expressed in tumor-associated ECs, including gliomas, ovarian tumor, breast tumor, and prostate cancer, suggesting the pro-angiogenic role of endothelial PDGFRs in disease.^30^ We show that adipose ECs exhibit consistently ∼4000 PDGFRα per cell, and the endothelial PDGFRβ increased in HFD-fed female mice, but not in male mice. These changes suggest potential synergistic mechanisms involving PDGFRs in female endothelial cells in the regulation of adipose angiogenesis and expansion.

In conclusion, our findings suggest that female mice preserve adipose tissue vascular health during HFD feeding by differentially regulating angiogenic signaling ligands and receptors. This sex-specific angiogenic regulation, including sustained high levels of VEGFR1 and PDGFRβ on endothelial cells, elevated VEGF-A, and higher baseline levels of PlGF, PDGF-AA, and PDGF-AB in adipose tissue, offers potential therapeutic strategies for obesity and metabolic disorders. Our findings are particularly relevant for bioengineering applications targeting vascularization and healthy adipose tissue expansion. Our study emphasizes the importance of studying sex as a biological variable, establishing its significance in angiogenic mechanisms related to obesity and metabolic health. Finally, understanding how VEGF signaling, PDGF signaling, and cross-family VEGF-PDGF interactions are regulated during healthy and unhealthy obesity in different sexes is important for advancing our knowledge and developing targeted interventions. These quantitative data on VEGF and PDGF ligands and receptors support systems biology approaches, enabling the development of computational models to delineate the complex regulation of angiogenic signaling in adipose tissue.

## METHODS AND MATERIALS

### Animals

All experiments in this study were reviewed and approved by the University of Washington Institutional Animal Care and Use Committee (IACUC). Male and female C57BL/6J mice were acquired from Jackson Laboratory at 6 weeks of age. All animals were maintained on a 14/10 light/dark cycle at 21°C and given access to food and water *ad libitum*. Upon receipt, the mice were randomly assigned to one of two diet groups. The high-fat diet group (HFD) received a diet consisting of 60% of calories from fat (Research Diets, D124921), while the low-fat diet group (LFD) served as the control and received a diet consisting of 10% of calories from fat (Research Diets, D12450Ji). After 12 weeks of respective diet treatments, five mice from each sex-diet group were euthanized with CO_2_. Gonadal white AT (gWAT) were processed or snap froze immediately for Week 12 measurements. The remaining HFD mice were randomly assigned to one of two diet groups. One group continued to receive the same 60% high-fat diet for long-term weight gain, while the other group switched from HFD to LFD for weight loss. The remaining LFD mice were kept on a 10% fat diet as a control. After another 12 weeks, gWAT samples were collected (processed or snap froze immediately) for Week 24 measurements.

### Stromal-vascular fraction (SVF) single-cell suspension preparation

Gonadal WAT was weighed and minced into small pieces (<1 mm^3) using fine surgical scissors. Approximately 0.5-1.0 g of minced tissue was then placed into gentleMACS C Tubes containing 5 mL of digestion buffer (Miltenyi Biotec, 130-105-808). The tubes were attached to the gentleMACS^TM^ Octo Dissociator (Miltenyi Biotec, 130-096-427). The built-in “mr_adipose_01” program was used for mechanical dissociation. The digested tissue was then filtered through a 70-μm cell strainer to remove undigested tissue and debris. The filtrate was collected and centrifuged at 300 × g for 10 minutes to separate the SVF from the floating mature adipocytes. The SVF cells were resuspended in cold stain buffer (Ca^2+^/Mg^2+^-free PBS + 0.5% BSA + 0.1% sodium azide, PH 7.4) at ∼ two million cells/mL.

### Flow cytometric measurements of VEGFRs and PDGFRs on the surface of SVF cells

SVF cells were aliquoted into 20 µL/FACS tubes and stained with APC-conjugated F4/80, APC-Cy7-conjugated CD31, FITC-conjugated CD45 antibodies per the manufacturer’s recommendation (5 µL /test, BioLegend). Phycoerythrin (PE)-conjugated VEGFR1 (R&D Systems), VEGFR2 (BioLegend), VEGFR3 (R&D Systems), PDGFRα (BioLegend), or PDGFRß (BioLegend) was added to each sample at the saturating concentrations^37,47^. Samples were incubated in the dark for 40Lminutes at 4 °C, washed twice with 2 mL of cold stain buffer, centrifuged at 400 g at 4 °C for 5 min, and re-suspended in 100 µl of stain buffer. To assess cell integrity, 2 µl of Sytox Blue was finally added to each tube as a liquid drop-in. Corresponding fluorescence-minus-one (FMO) controls were used for the evaluation of the nonspecific binding of monoclonal antibodies to identify the positive/negative boundary for each fluorophore expression. The FMO control contained all the fluorophores in a panel, except for the one being measured. Flow cytometry was performed on a Beckman Coulter CytoFLEX S Flow Cytometer. Samples were vortexed immediately prior to placement in the flow cytometer. Prior to sample acquisition, PE voltage settings were finalized and Quantibrite^TM^ PE beads (BD, Cat. No. 340495) were collected to establish PE calibration curves^54^. The arbitrary PE fluorescence intensities were converted to the number of PE-labelled receptors per cell using Quantibrite^TM^ PE beads, as previously described.^38,47,53–55^ Flow cytometric data analysis was performed using Kaluza analytical software.

### ELISA measurements of VEGF and PDGF ligands in gonadal white AT

Snap-frozen gWAT sample (∼0.3 g/sample) was added to a gentleMACS M Tube containing 3 mL cold homogenization buffer that consists of 95% 1X Cell Lysis Buffer (Cell Signaling, 9803), 1% Antifoam L-30 Emulsion (Sigma-Aldrich, STS0100), and 4% 25X EDTA-free Protease Inhibitor Cocktail (cOmplete™, 11873580001). The M tube was then attached to the gentleMACS^TM^ Octo Dissociator (Miltenyi Biotec, 130-096-427). The built-in “ Protein_01” program was used for tissue homogenization. The homogenized tissue was centrifuged at 400 × g at 4°C for 5 minutes to separate the cell pellets from the floating lipid cake layer. The cells pellets and lipid-free supernatant were transferred into 2-mL Eppendorf tubes. The cell lyses were centrifuged at 14,000 ×g at 4°C for 10 minutes to let the cell pellets release proteins. The protein-containing supernatant to new tubes, stored at -20C, or used immediately. The total protein concentration of each sample was determined using Pierce™ BCA Protein Assay Kits (23225). The concentrations of VEGF-A, PlGF, and PDGFs were measured using Invitrogen ELISA Kits: (1) Mouse VEGF-A ELISA Kit (BMS619-2), (2) Mouse PDGF-AA ELISA Kit (EM61RB), (3) Mouse PDGF-BB ELISA Kit (EM63RB), (4) Mouse PDGF-AB ELISA kit (EM62RB), and (5) Mouse PlGF-2 ELISA Kit (EMPGF). The ligand concentrations were normalized to the total protein concentration of each gWAT sample.

### Statistical analysis

Measurements of body weights, tissue weights, ligand concentrations, and receptor numbers were shown as mean ± standard error of the mean of four to nine mice. Statistically, differences in the measurements between two different sex-diet groups were determined by a One-Way ANOVA with Tukey post-hoc analysis (* p<0.05, ** p<0.01, *** p<0.001, **** p< 0.0001) using Originlab Pro software (2022).

### Single-nucleus RNA Sequencing Sample Preparation and Nuclei Isolation

Gonadal white adipose tissue (gWAT) samples were snap-frozen in liquid nitrogen and stored at - 80°C until processing. snRNA-seq data are from one mouse from each of the six groups: Week 12 male HFD and LFD, Week 24 male HFD and LFD, and Week 24 female HFD and LFD. Nuclei were isolated using the Chromium Nuclei Isolation Kit with RNase Inhibitor (PN-1000494, 10x Genomics), following the manufacturer’s protocol. Briefly, tissue was thawed on ice, minced, and homogenized in the lysis buffer provided in the kit. The homogenate was filtered and centrifuged at 500 × g for 10 minutes at 4°C to collect nuclei. The isolated nuclei were washed, resuspended in buffer, and assessed for integrity and concentration using trypan blue exclusion and a hemocytometer.

### Probe Hybridization and GEM Generation

Isolated nuclei were fixed and processed using the Chromium Next GEM Single Cell Fixed RNA Sample Preparation Kit (PN-1000414), Chromium Next GEM Chip Q Single Cell Kit (PN-1000422), and Chromium Fixed RNA Kit, Mouse Transcriptome (PN-1000496) according to the protocol outlined in the Chromium Fixed RNA Profiling User Guide (CG000527, Rev E). Fixed nuclei were hybridized with whole transcriptome probes overnight at 42°C. Following hybridization, nuclei were pooled and loaded into a Chromium Next GEM Chip Q, along with Gel Beads and reagents, for generation of Gel Beads-in-Emulsion (GEMs). The barcoding process incorporated unique GEM and probe barcodes, enabling sample multiplexing and demultiplexing during data analysis.

### Library Construction and Sequencing

Barcoded RNA fragments were recovered from the GEMs, pre-amplified, and subjected to library construction using the Chromium Fixed RNA – Gene Expression Library Kit. Sample Index PCR was performed to generate final libraries, which were quantified using an Agilent Bioanalyzer. Libraries were sequenced on an Illumina NovaSeq 6000 platform.

### Data Processing and Analysis

Sequencing data were processed using the Cell Ranger pipeline (10x Genomics), aligning reads to the mouse genome reference. We processed snRNA-seq FASTQ reads by aligning them to the human reference genome (GRCh38) using the 10X Genomics Cell Ranger pipeline (v.9.0.0)^56^ to generate gene-barcode matrices. Low-quality nuclei were filtered out based on three criteria: a mitochondrial read proportion exceeding 15%, fewer than 6,000 detected features, or fewer than 15,000 total counts. The remaining nuclei were clustered, normalized, and scaled using Seurat (v.5.0)^57^. Principal Component Analysis (PCA) was performed on the processed data, retaining 50 principal components. To integrate multiple samples and correct for batch effects, the *RunHarmony()* function from the Harmony R package (v1.2.3)^58^ was applied to the PCA dimensions. The harmonized principal components were then used as input for UMAP embedding with Seurat.

For cell type annotation, we used the *FindAllMarkers()* function to identify differentially expressed genes (DEGs) across clusters. DEGs were used to assign cell type identities by referencing previously reported annotations in the literature^59,60^.For differential expression analysis between experimental groups, we analyzed genes that were present in at least 25% of nuclei in both conditions. DEGs were identified using Seurat’s *FindMarkers()* function with the Wilcoxon rank-sum test.

## Data availability statement

All data are included in the main text and supplementary files. Other raw data are available upon reasonable request.

## Code availability statement

No custom code or mathematical algorithms were used in this study.

## Acknowledgments

This research used statistical consulting resources provided by the Center for Statistics and the Social Sciences, University of Washington. We also thank Dr. Ana Valencia from the UW Medicine Division of Metabolism, Endocrinology, and Nutrition, for her support with regular mouse weighing and monitoring. This work was supported by the National Science Foundation (Grant No. 1653925), the National Institute of Heart, Lung, and Blood (5R01HL159946-04) and by the Saint Louis University Startup Fund (VC). The content is solely the responsibility of the authors and does not necessarily represent the official views of the funding agencies.

