## Supplementary figures and images for "Sex Differences in VEGF and PDGF Ligand and Receptor Protein Expression during Adipose Tissue Expansion and Regression"

### Suppl Fig 1.tif

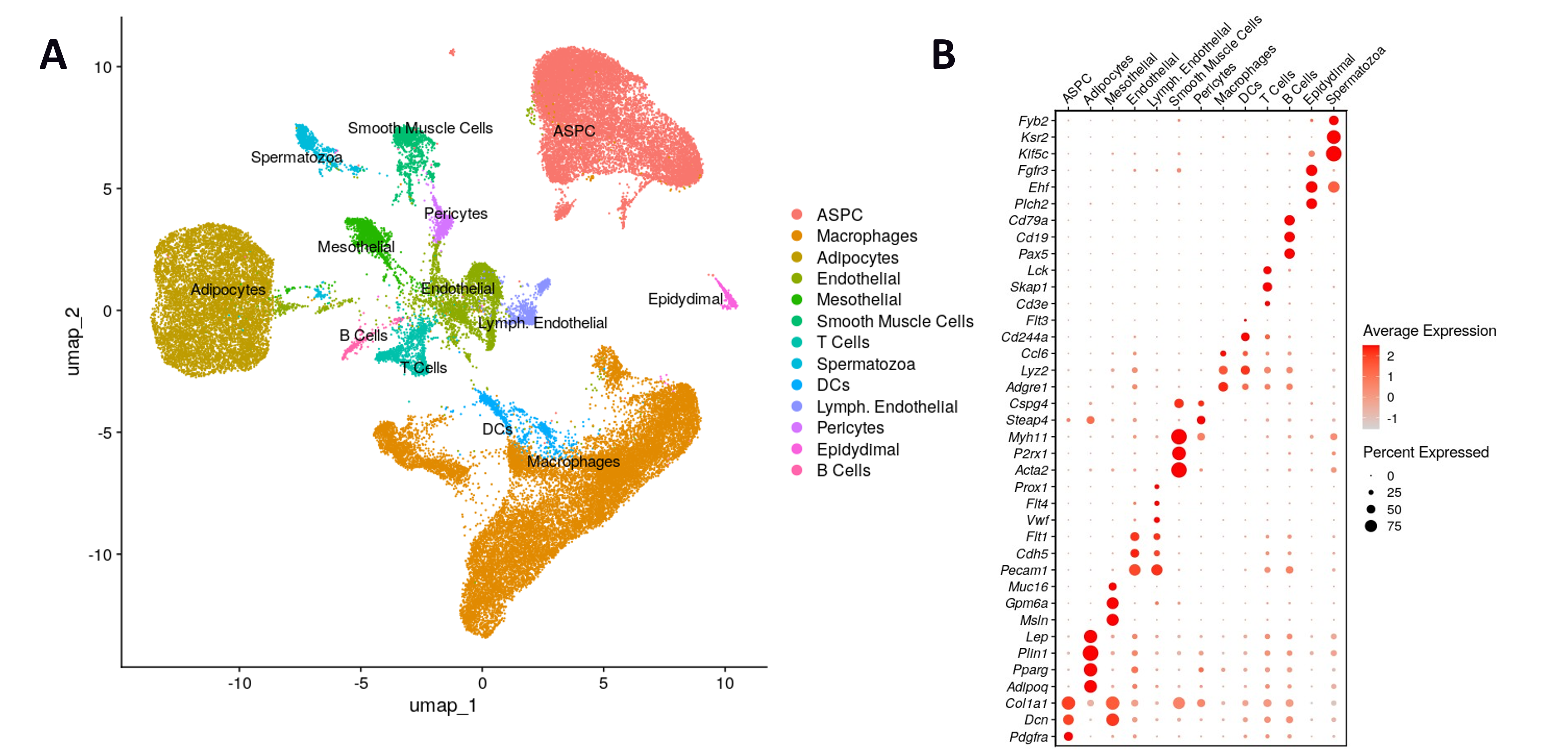

### Suppl Fig 2.tif

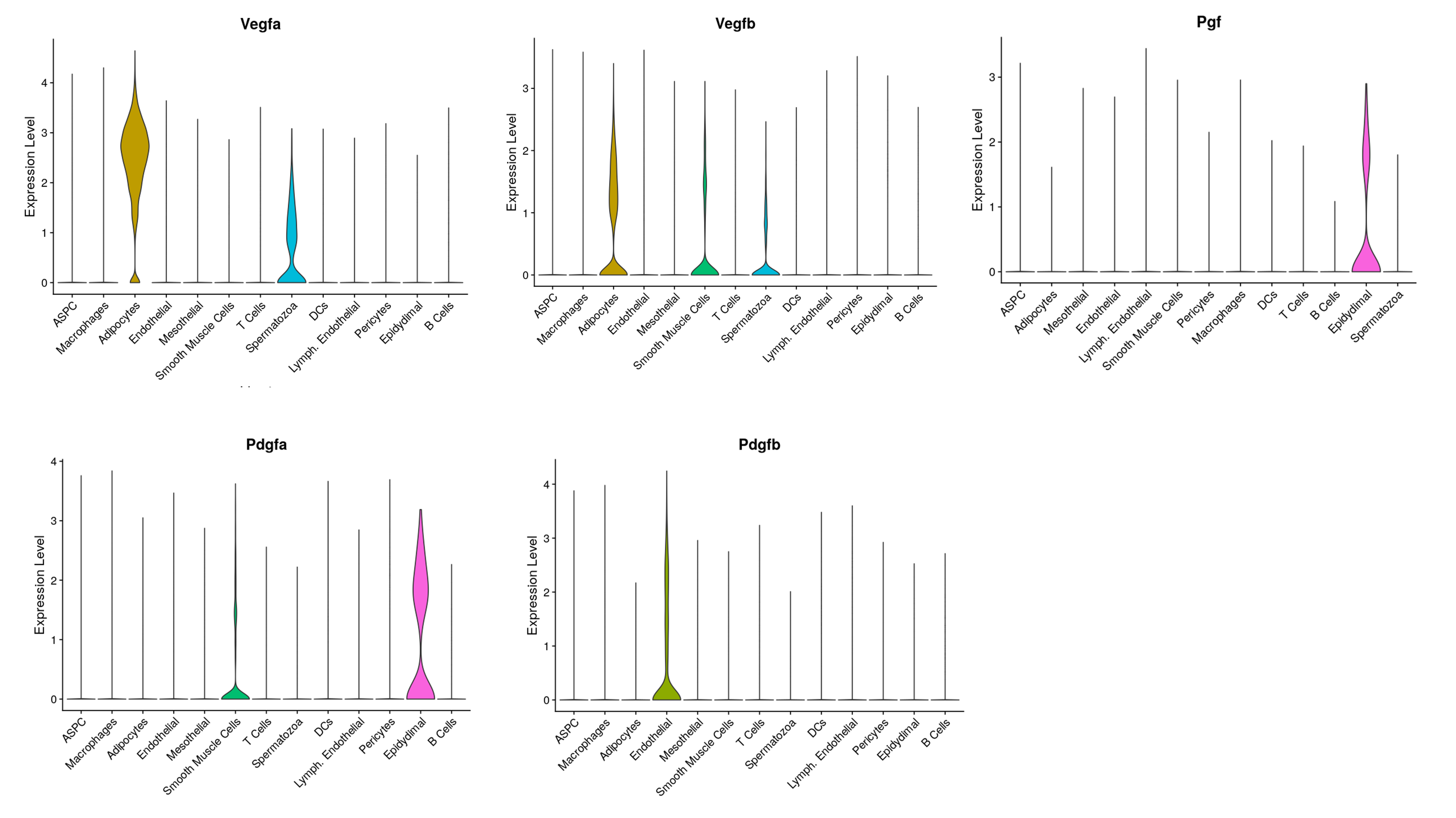

### Suppl Fig 3.tif

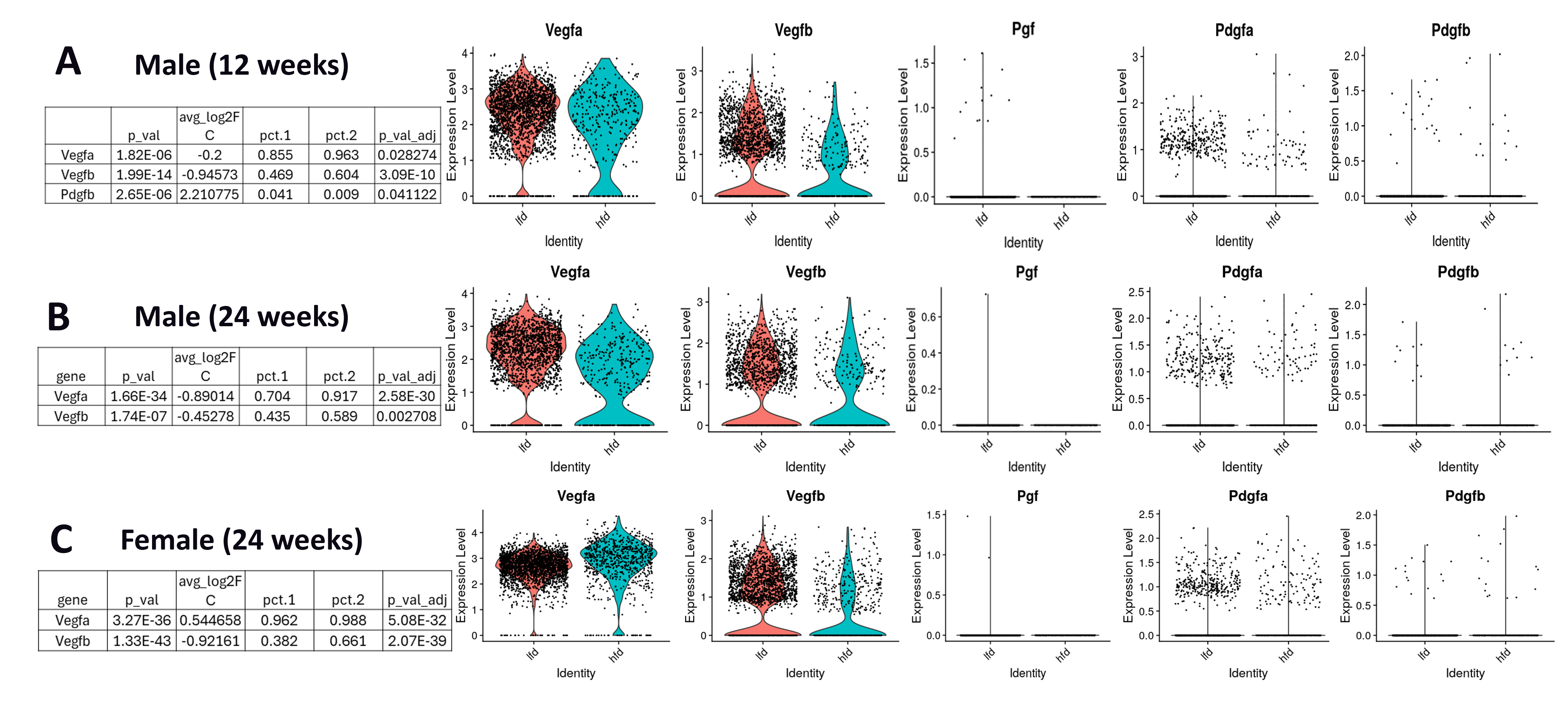

### Suppl Fig 4.tif

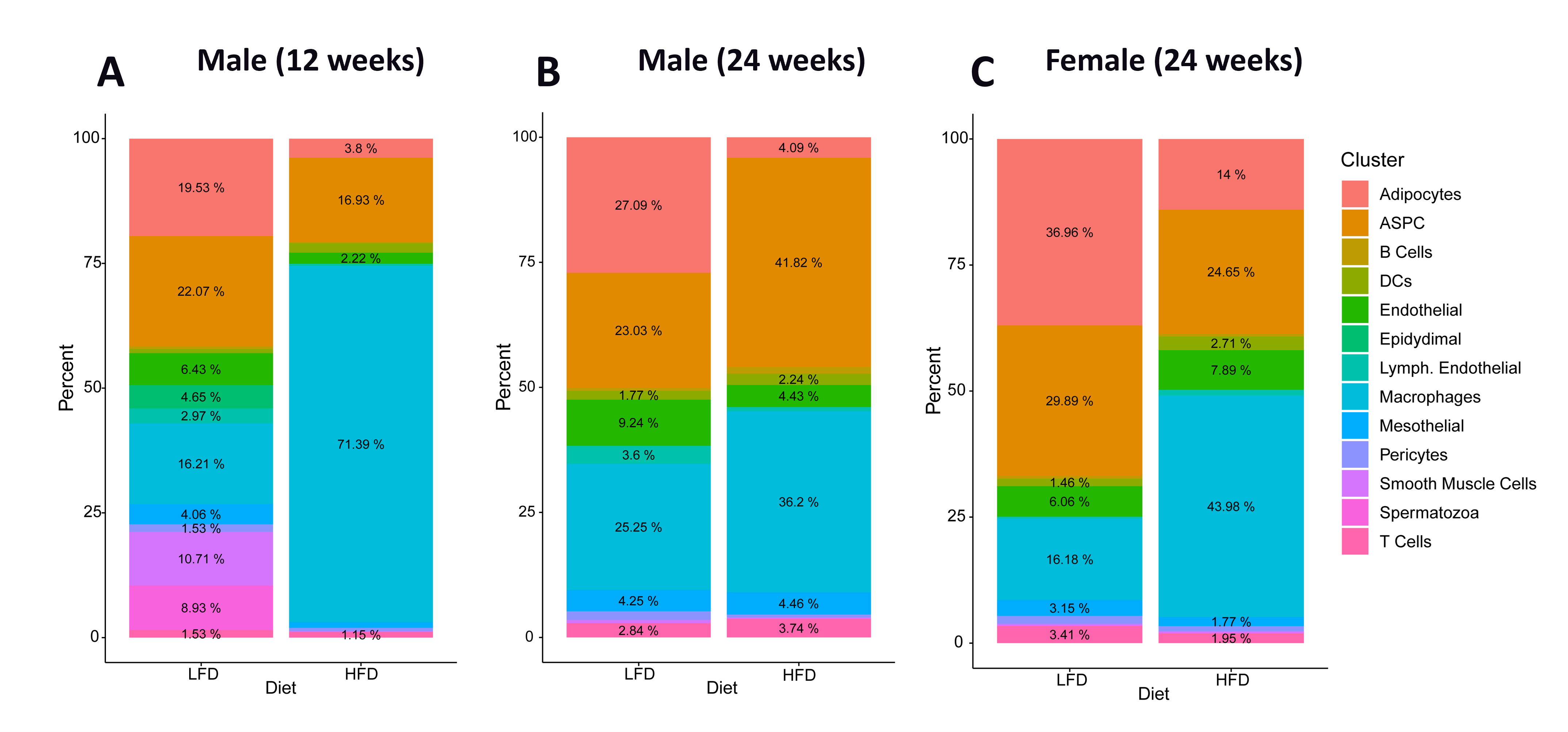

### Suppl Fig 5.tif

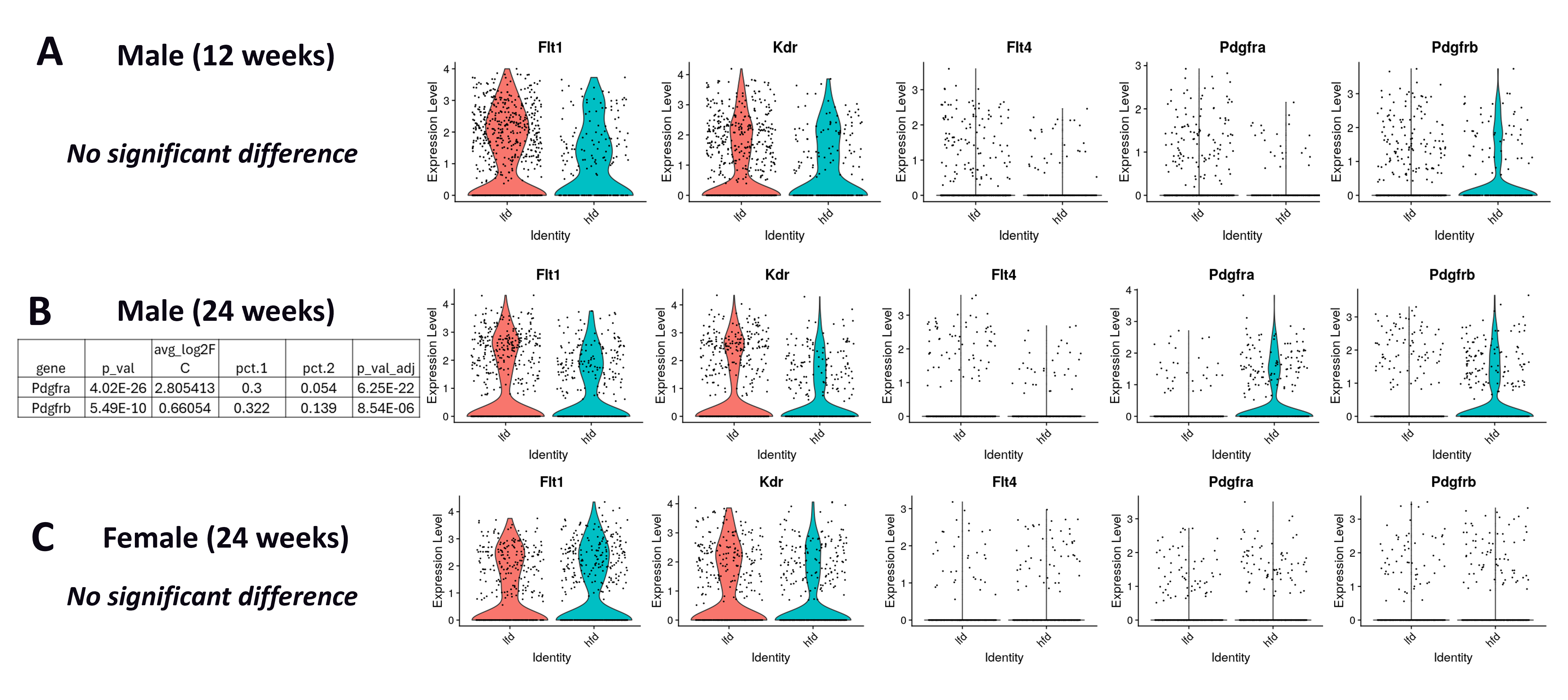

### Suppl Fig 6.tif

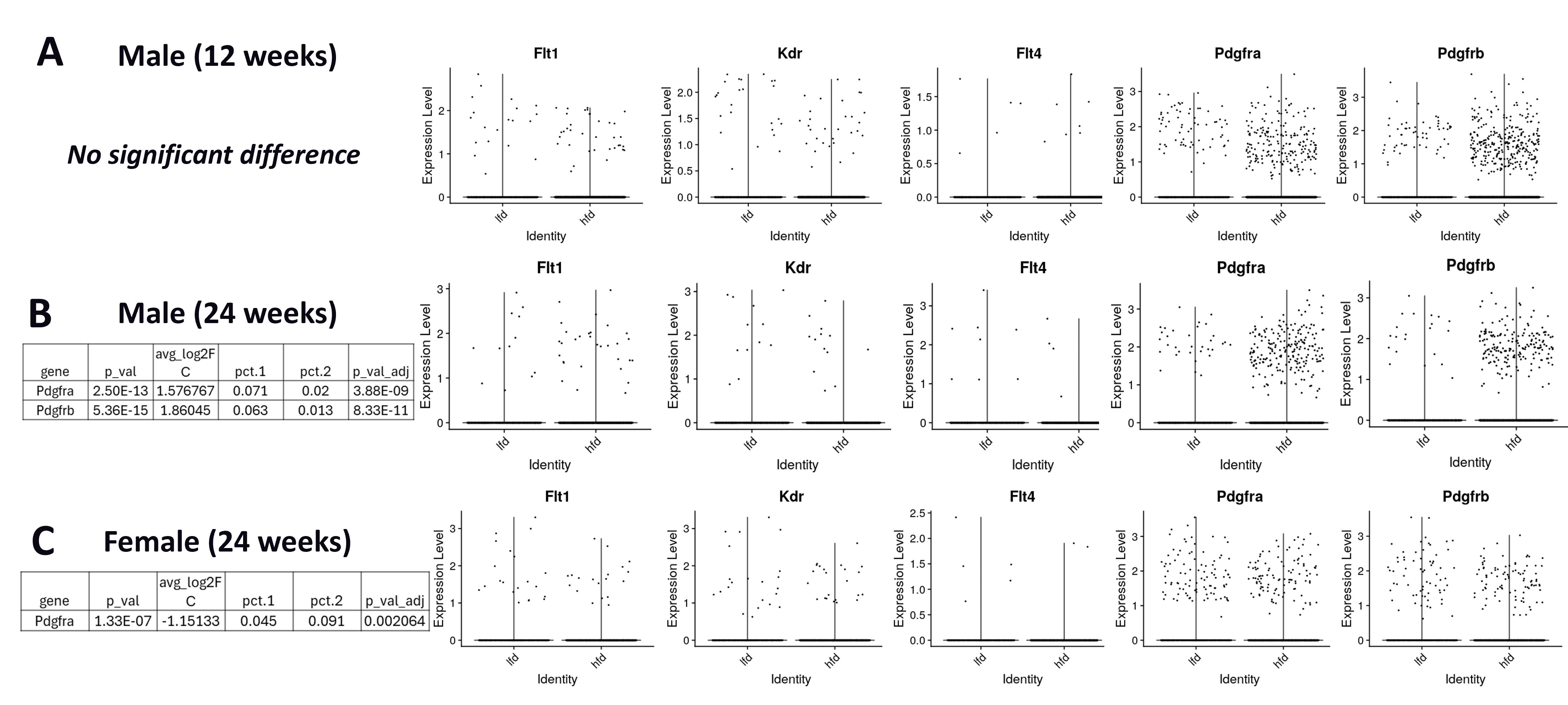
